# A meta-analysis of periodic and aperiodic M/EEG components in Parkinson’s disease

**DOI:** 10.1101/2025.07.07.663454

**Authors:** Hamzeh Norouzi, Magdalena Ietswaart, Jason Adair, Gemma Learmonth

## Abstract

Parkinson’s disease is characterised by a range of motor and non-motor changes that can negatively impact quality of life. Many studies have identified potential clinical electrophysiological biomarkers of Parkinson’s disease with an aim of developing new methods of identifying at-risk patients, and to form the basis of therapeutic interventions. However, these studies do not present consistent results, and a formal meta-analysis is warranted to identify reliable M/EEG characteristics across datasets. In this meta-(re-)analysis of open-access M/EEG datasets (n = 6; 4 EEG and 2 MEG), we compared periodic and aperiodic characteristics of resting-state recordings in 368 patients with Parkinson’s disease and 570 age-matched healthy controls. Specifically, we compared the power and peak frequency of the aperiodic-adjusted alpha and beta oscillations, and the aperiodic exponent and offset across the two groups. Using spectral parametrisation, individuals with Parkinson’s disease had higher alpha-band power and a slower alpha peak frequency compared to controls, however no group differences in beta-band power and peak frequency were identified in this resting state data. Parkinson’s patients were furthermore found to have consistently higher aperiodic offset and exponent, possibly indicative of increased cortical inhibition. In conclusion, this large cohort meta-analysis points to a broadly consistent pattern of both periodic and aperiodic changes in Parkinson’s patients in M/EEG signal that may be used to develop diagnostics and targeted interventions in the future.

## 1. Introduction

Parkinson’s disease (PD) is a progressive neurodegenerative disorder caused by a loss of dopamine-producing neurons in the substantia nigra, which plays a critical role in motor control. PD primarily affects the execution of smooth movement, and is characterised by tremors, rigidity, bradykinesia and postural and gait impairments (Vinding et al. 2024a). Yet PD is also a multi-faceted disorder and is often accompanied by a range of non-motor symptoms, including cognitive decline, fatigue, apathy, visuospatial disturbances and sleep disruption (Kalia and Lang 2015; Vinding et al. 2024a; Yassine et al. 2023).

Electrophysiological recordings have identified specific changes in cortical oscillatory activity in patients with Parkinson’s disease, particularly within the theta (4-7 Hz), beta (13-30 Hz) and gamma (40+ Hz) bands (Foffani and Alegre 2022). One of the most consistent markers of “oscillopathy” in PD is the beta activity associated with motor preparation and execution. During active movement in healthy individuals, beta power gradually desynchronises in anticipation of an action, reaches its lowest point during movement execution, and then rapidly rebounds to a level above baseline after the movement ends (Szul et al. 2023). In contrast, this pattern is disrupted in PD patients, with the beta desynchronisation before and during movement execution diminished, and the post-movement beta rebound also reduced in magnitude (Boon et al., 2019; Heinrichs-Graham, Wilson, et al., 2014; Mustile et al., 2023).

Tonically high levels of beta have also been observed at rest in Parkinson’s disease and these elevated periods of beta have been directly associated with motor dysfunction in the form of bradykinesia (Zavala et al. 2015) and freezing of gait (Toledo et al. 2014). This has been corroborated by invasive electrophysiology which has identified elevated resting-state beta activity within the basal ganglia nuclei and cortex (Eisinger et al. 2020; Thomas Binns et al. 2025; Tinkhauser, Pogosyan, Little, et al. 2017a; Tinkhauser, Pogosyan, Tan, et al. 2017). However, the literature is mixed, with non-invasive resting state studies variously describing higher beta activity in PD patients relative to controls (Cao et al. 2020; McKeown, Jones, et al. 2024a), lower beta activity (Gimenez-Aparisi et al. 2023; Jaramillo-Jimenez et al. 2023a; Olde Dubbelink et al. 2013), and no difference between patients and controls (Vinding et al. 2024b; Wang et al. 2022a; Wiesman, Madge, et al. 2025a). Further, dopaminergic medication appears to effectively suppress beta activity at the cortical and subcortical levels (Binns et al. 2025). Thus, although beta oscillations may show some promise as a potential biomarker to dissociate PD from controls, this potential is limited due to its sensitivity to dynamic physiological and pharmacological conditions.

In contrast, the lower oscillatory frequencies, particularly within the alpha band (8-12 Hz) have been explored less often in Parkinson’s disease. In healthy adults, alpha rhythms typically reflect decreased excitability of the underlying cortex and are modulated by both tonic and selective attention (Jensen 2024; Thut et al. 2006). Several studies have reported an increase in alpha power and decreased peak frequency in PD, which appear to be positively associated with non-motor disease symptoms, specifically cognitive decline (Anjum et al. 2024; Olde Dubbelink et al. 2013; Özkurt 2024; Sinanovic et al. 2005; Wiesman et al. 2023a; Wiesman, Madge, et al. 2025b; Yassine et al. 2023; Ye et al. 2022). However, this pattern is also inconsistent, with others reporting lower alpha, or no differences between patients and healthy controls (Bosboom et al. 2006; George et al. 2013; Jaramillo-Jimenez et al. 2023b; Mikkel Vinding et al. 2022).

Electrophysiological signals reflect a combination of periodic (oscillatory) and aperiodic components, which are postulated to arise from distinct neural mechanisms (Donoghue et al. 2020; Hughes et al. 2012; Wen and Liu 2016). The aperiodic component of electrophysiological signals carries potential physiological significance and exhibits dynamic modulation in response to factors such as age, cognitive state, and task demands (Donoghue et al. 2020). The aperiodic component can be characterized by two parameters; the exponent is equivalent to the negative slope of the power spectrum, and the offset reflects the uniform shift of power across frequencies (Donoghue et al. 2020; Hughes et al. 2012; Wen and Liu 2016). The aperiodic exponent has been linked to the balance and integration of underlying excitatory and inhibitory synaptic currents (E/I ratio) (Martínez-Cañada et al. 2023; van Nifterick et al. 2023), while changes in the offset reflect shifts in the broadband signal, particularly at lower frequencies (Manning et al. 2009; Miller et al. 2014).

Recent advancements in electrophysiological signal analysis have made it possible to detect changes in the excitation-inhibition ratio, which may be detectable during (and potentially in the prodrome of) disease processes. A link between the aperiodic exponent and the E/I ratio in the brain is frequently reported (Martínez-Cañada et al. 2023; van Nifterick et al. 2023). Ketamine, by blocking NMDA receptor-mediated input to inhibitory interneurons, induces cortical disinhibition and increases glutamate release (Grunze et al. 1996; Maccaferri and Dingledine 2002). In contrast, thiopental, a GABA_A_ receptor modulator, suppresses cortical excitation (Pittson et al. 2004). Both drugs have been shown to modulate the aperiodic exponent in healthy individuals (Cortes-Briones et al. 2025). However, other pharmacological manipulations of inhibitory neural mechanisms have failed to support the hypothesis that the aperiodic exponent reliably reflects the E/I ratio in electrophysiological activity (Salvatore et al. 2024).

Both non-invasive and invasive electrophysiology in humans and animals have identified increased synchronised neural activity in PD patients (reflected by greater offset) as well as elevated spontaneous neuronal activity in the basal ganglia and in the cortical level, which shifts the balance of the normal E/I ratio (indicated by larger exponent, i.e. steeper slope in the power spectrum) in the brain (Donoghue 2024; Filion and Tremblay 1991; Helson et al. 2023; Rosenblum et al. 2023; Wang et al. 2022b; Wiest et al. 2023). Such increases in the aperiodic exponent and offset in PD patients relative to controls have been documented (McKeown, Jones, et al. 2024b; Vinding et al. 2024a; Wiesman et al. 2023a). However, several studies have reported either a reversed pattern (Mostile et al. 2019; Peng et al. 2024; Wu et al. 2023) or no significant differences in aperiodic offset and exponent between individuals with Parkinson’s disease and healthy controls (Bernasconi et al. 2023a; Clark et al. 2023; Martin et al. 2018; Monchy et al. 2023; Pardo-Valencia et al. 2024).

A recent advancement in spectral parameterisation of M/EEG data has highlighted that previous studies of group differences in periodic oscillatory power may be confounded by changes in aperiodic electrophysiological components (Donoghue et al. 2020; Kopčanová et al. 2024). By first extracting the offset and exponent separately from periodic components, we aim to identify more subtle changes in cortical activity that may point to more consistent and reliable biomarkers of PD that have previously been obscured.

Collectively, these findings suggest the need to identify consistent patterns of altered cortical electrophysiology associated with Parkinson’s disease to develop novel biomarkers to diagnose PD, predict its onset, and to formulate more effective treatments. In this study, we leverage large-scale open-source M/EEG resting-state datasets in patients with Parkinson’s disease and healthy controls. We applied a single, custom processing pipeline to perform a spectral parameterisation re-analysis of the raw datasets, followed by 6 meta-analyses to synthesise the evidence for between-group differences in the power and peak frequency of aperiodic-adjusted alpha and beta oscillations, and the aperiodic exponent and offset between PD patients and healthy controls.

## 2. Methods

### 2.1. Study Selection

Studies were identified through electronic databases, including bioRxiv, medRxiv, PsyArXiv, PsycINFO, PubMed, ScienceDirect, Scopus, and Web of Science from inception until September 2024. The search terms used to identify the datasets were (“Parkinsons’s disease” OR “Parkinsons” OR “Parkinsonian” OR “Parkinson”) AND (“resting state” OR “resting-state” OR “rest state” OR “rest-state” OR “rest”) AND (“EEG” OR “MEG” OR “electroencephalogram” OR “magnetoencephalogram”). We included datasets where raw resting-state EEG or MEG was available from both human Parkinsonian patients and healthy controls. A total of 226 studies were identified (Figure 1). After removing duplicates, 159 unique records were screened and 123 records were excluded for the following reasons: did not include electrophysiological signal recordings, described animal models, recorded deeper brain structures using ECOG, LFP, or DBS, and investigated mild traumatic brain injury, multiple sclerosis, or Parkinson’s disease patients during sleep. Of the 36 remaining records, 13 were further excluded because they: only included patients with no control group, did not include resting-state data (Agouram et al. 2024; Bernasconi et al. 2023b; Bringas et al. 2020; Cockburn et al. 2014; Lee et al. 2023; Pellegrini et al. 2024; Rassoulou et al. 2024; Singh et al. 2018; Stuart et al. 2019; Wang et al. 2022c; Xefteris et al. 2020; Zhang et al. 2022a; Zimmermann et al. 2015), and 17 records were not retrieved (did not allow data sharing or did not respond to our data-sharing requests). Ultimately, resting-state data from six publicly available studies were included: San Diego (Rockhill et al. 2020), Turku (Railo, 2021), New Mexico (Cavanagh, 2021), and Iowa (Singh et al. 2023) and 2 were shared with us upon request: OMEGA (Niso et al. 2016) (the original report from OMEGA dataset that we refer to is (Wiesman et al. 2023a), and NatMEG (Vinding et al. 2023a). Details of the searches are illustrated in Figure 1.

**Figure 1.**
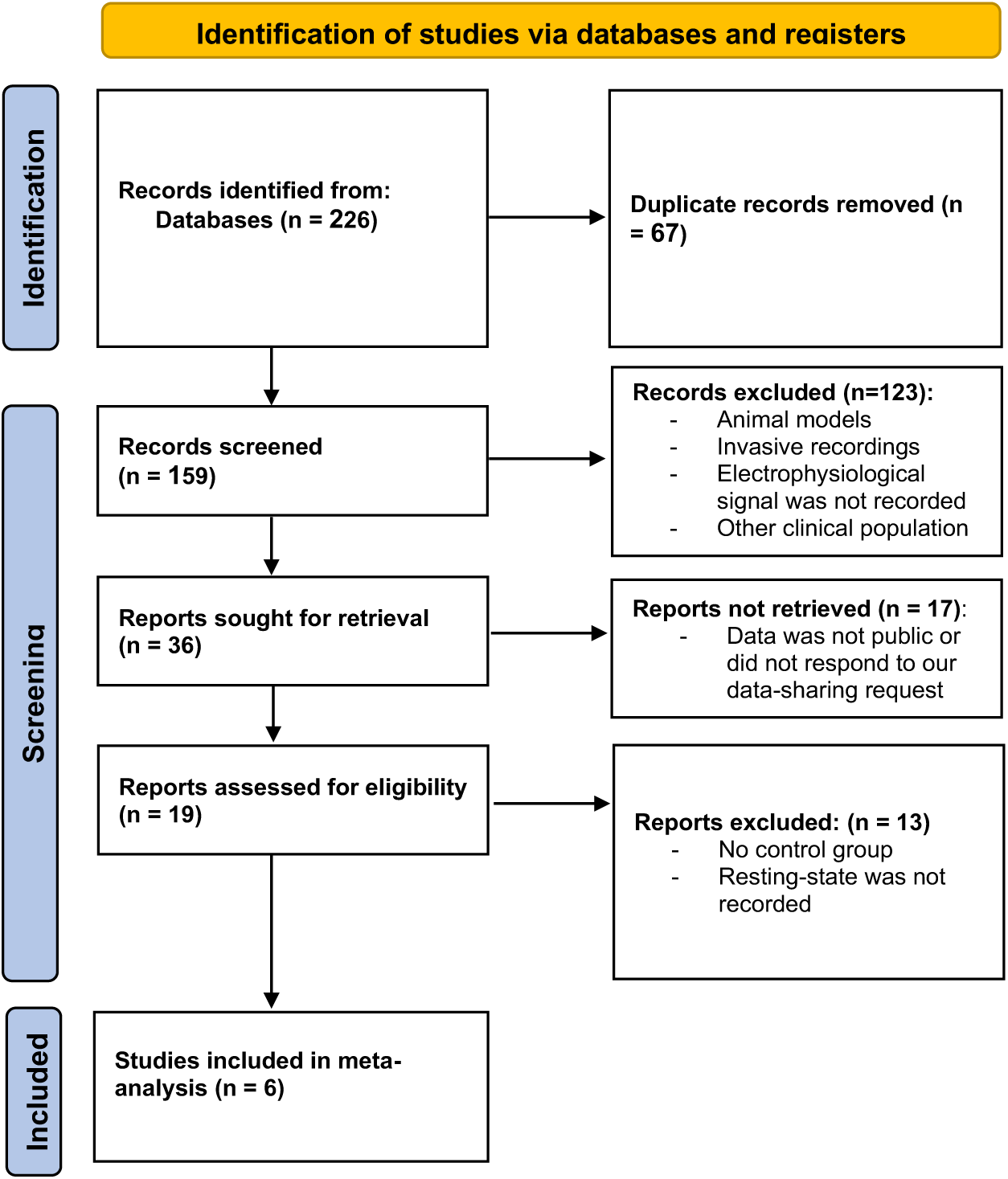
Flow diagram documenting the electronic database searches, abstract screening, and the reasons for excluding studies. Adapted from Preferred Reporting Items for Systematic Reviews and Meta-Analyses (PRISMA; Page et al., 2021).

The datasets entered into the pre-processing stage involved a total of 1092 participants, of which 426 participants were PD patients, and 666 were healthy aged-matched control individuals. The San Diego, Turku, Iowa, and New Mexico datasets were recorded using EEG with various numbers of electrodes, and the OMEGA and NatMEG datasets were recorded using MEG. The San Diego, Turku, and New Mexico datasets recruited patients both in ON and OFF medication states, however, we only included recordings for the ON medication state in this analysis. The characteristics of each dataset are presented in Table 1 and are described below.

**Table 1.** Characteristics of data cohorts assessing resting-state EEG/MEG in Parkinson’s patients and healthy controls.

| DATASET | Total participants | Participants included in meta-analysis | M/EEG | Resting state recording duration (min) | PD Age | PD Sex | Mean PD duration (years) | Mean H&Y | Mean UPDRS score (ON medication) | HC Age | HC Sex | SpecParam analysis performed |
| --- | --- | --- | --- | --- | --- | --- | --- | --- | --- | --- | --- | --- |
| San Diego | 16 PD, 16 HC | 13 PD, 10 HC | EEG | > 3 | 62.6 (8.3) | 8F | 4.9 (3.6) | - | 33.7<sup>‡</sup> (10.9) | 63.5 (9.6) | 9F | No |
| Turku | 20 PD, 20 HC | 6 PD, 20 HC | EEG | > 2 | 69.5 (7.7) | 10F | 6 (5.0) | - | 23.0* (10.2) | 67 (6.1) | 12F | No |
| Iowa | 132 PD, 49 HC | 115 PD, 42 HC | EEG | > 2 | 68.1 (8.0) | 43F | 4.1 (3.5) | 1.45 | 12.5* (7.2) | 70.9 (7.6) | 23F | No |
| New Mexico | 27 PD, 25 HC | 24 PD, 23 HC | EEG | ≈ 9 | 69.7 (8.7) | 9F | 5.4 (4.1) | - | 23.4* (9.9) | 69.3 (9.6) | 9F | Yes |
| OMEGA | 166 PD, 488 HC | 154 PD, 411 HC | MEG | ≥ 5 | 65.0 (9.1) | 42F | 6.8 (4.3) | 1.96 (0.7) | 32.7* (14.8) | 44.4 (20.8) | 305F | Yes |
| NatMEG | 66 PD, 68 HC | 56 PD, 64 HC | MEG | ≥ 3 | 64.4 (9.8) | 22F | 4.0 (3.4) | 1.63 (0.77) | 19.3<sup>‡</sup> (11.1) | 63.3 (8.7) | 31F | Yes |

(<sup>‡</sup>) UPDRS overall score, (\*) UPDRS-III scores. PD = Parkinson's disease, HC = Healthy controls, F = females, UPDRS = Unified Parkinson's Disease Rating Scale, H&Y = Hoehn and Yahr Score.

#### Dataset 1: San Diego, USA (Rockhill et al. 2020)

16 PD patients (8 women, mean age = 62.6 years, SD = 8.3) and 16 healthy controls (9 women, mean age = 63.5, SD = 9.6) were recorded. All patients had mild to moderate PD severity (Hoehn and Yahr stage II and III). PD patients were recorded both ON and OFF medication. EEG was recorded for at least 3 minutes using a 32 channel ActiveTwo (Biosemi Instrumentation system) using a sampling rate of 512Hz while participants kept their eyes open and maintained fixation.

#### Dataset 2: Turku, Finland (Railo, 2021)

Twenty PD patients (10 women, mean age = 69.5, SD = 7.7) and twenty healthy controls (12 women, mean age = 67, SD = 6.1). EEG was recorded for at least 2 minutes using 64 active electrodes with a sampling rate of 500Hz during eyes open and eyes closed. Twelve patients were recorded OFF medication, and five patients were recorded ON medication but only the eyes open, ON medication recordings were selected for the meta-analysis.

#### Dataset 3: Iowa, USA (Singh et al. 2023)

132 PD patients (43 women, mean age = 68.1, SD = 7.98) and 49 healthy controls (23 women, mean age = 70.91, SD = 7.62) were tested. 64 channel EEG was recorded with eyes open for at least 2 minutes with a sampling rate of 500Hz. All patients were tested ON medication.

#### Dataset 4: New Mexico, USA (Cavanagh, 2021)

25 PD patients (9 women, mean age = 69.68, SD = 8.73) and 25 age- and sex-matched controls (9 women, mean age = 69.32, SD = 9.58) were tested. EEG was recorded for around 9 minutes with 64 Ag/AgCl electrodes with a sampling rate of 500 Hz during both eyes closed and eyes open. PD patients were tested both ON and OFF medication. The eyes open, ON medication data was used in the meta-analysis.

#### Dataset 5: OMEGA, Canada (Niso et al. 2016)

166 PD patients (46 women, mean age = 64.95, SD = 9.12) and 488 healthy controls (307 women, mean age = 44.42, SD = 20.83) were tested. At least 5 minutes eyes open resting-state data was recorded using a 275-channel whole-head CTF MEG system at a sampling rate of 2400 Hz and with an antialiasing filter with a 600 Hz cut-off. (Note: The OMEGA dataset is continuously expanding, with new participants, both patients and healthy controls, are being regularly added. In the original report, the dataset comprised 79 individuals with Parkinson’s disease and 65 healthy controls (Wiesman et al. 2023a).

#### Dataset 6: NATMEG, Sweden (Vinding et al. 2023b)

66 PD patients (28 women, mean age = 65.6, SD = 9.5) and 68 healthy controls (27 women, mean age = 63.93, SD = 8.4) were tested. At least 3 minutes of resting-state data was recorded using a Neuromag TRIUX 306-channel MEG system, with 102 magnetometers, with 1000 Hz sampling rate, with an online 0.1 Hz high-pass filter and 330 Hz low-pass filter while the participants sat with their eyes closed. The average head movements were not significantly different between groups (t(115.8) = 0.55, *p* = 0.58).

### 2.2. Preprocessing

The OMEGA, NatMEG and New Mexico datasets had previously been subjected to spectral parameterisation (Table 2; reported in the respective source publications), but for the purposes of this study we developed a single processing pipeline in order to re-analyse the 6 datasets using a single set of common parameters. Each of the datasets were first re-sampled to a uniform sampling rate of 500 Hz, then filtered between 1-100 Hz using a zero phase Infinite Impulse Response (IIR) filter and detrended to remove slow drifts. Independent component analysis (ICA) was applied to the filtered signal and non-brain components were excluded automatically using MNE-ICALabel (Li et al. 2022) for the 4 EEG datasets, and empty-room signal was used to reduce environmental noise in the 2 MEG datasets. The reference channels were available for the OMEGA dataset and were used to identify non-brain components. Both gradiometer and magnetometer channels were available for the NatMEG dataset, and only magnetometers were available in the OMEGA dataset, therefore only magnetometer channels were used for analysis of both MEG datasets. Next, the pre-processed signal was segmented into 1 second epochs. Each dataset resulted in different number of epochs due to the variable length of the recordings. Due to poor signal quality or data corruption, 9 subjects from San Diego dataset, 23 subjects from Iowa, 5 subjects from New Mexico, 45 subjects from OMEGA, and 11 subjects from NatMEG were excluded. A total of 938 subjects (368 PD patients and 570 controls) were included in the final spectral parametrisation stage.

**Table 2.**
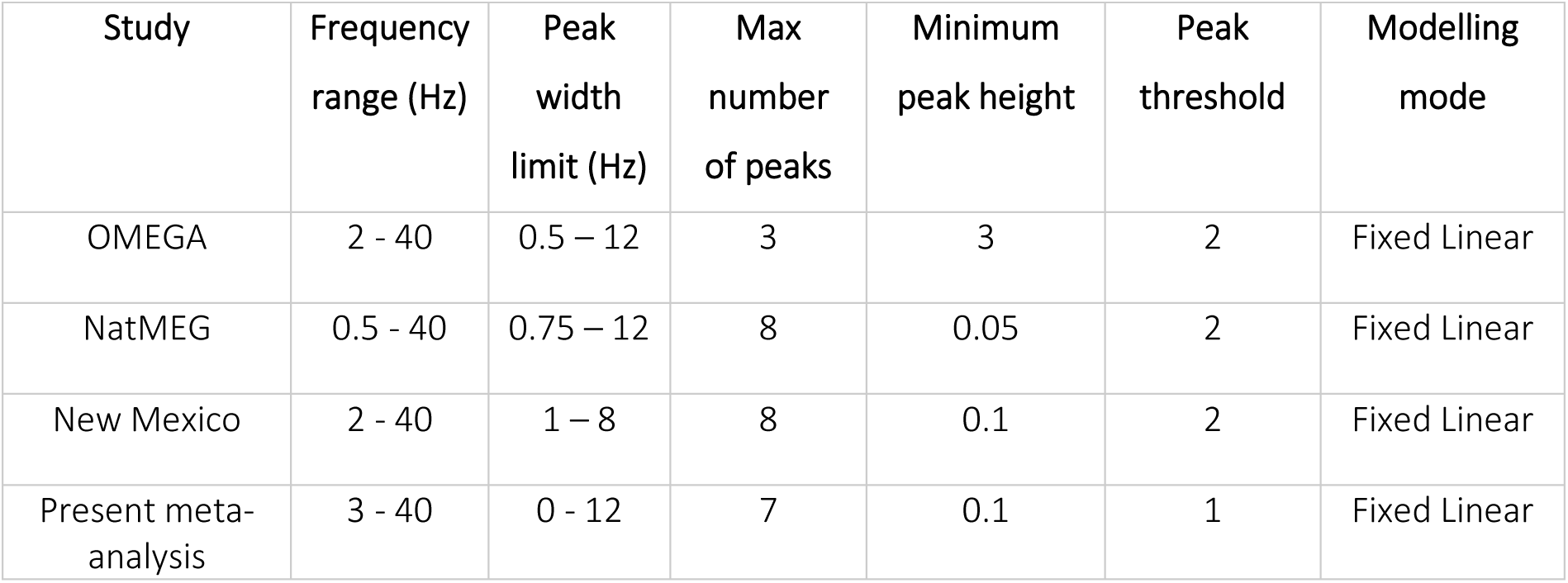
Comparison of SpecParam settings between previously parametrised datasets and the present meta-analysis.

### 2.3. Spectral parameterisation

The power spectrum of each epoch was estimated using Welch’s method with a segment length of 5000 samples (10 * sampling rate), 50% overlap, and a Hanning window. Power spectrum signal was then averaged over 3 different regions of interest involving the Frontal, Central, and Posterior electrodes, as well as an average across all electrodes, referred to as the AllScalp region. Next, the power spectrum of each region was parameterized using the *specparam* toolbox (Donoghue et al. 2020) in MNE python (Larson et al. 2024) using the following settings: peak width limits ([0, 12]), peak threshold (1), aperiodic mode (fixed), maximum number of peaks (7), and minimum peak height (0.1). Each of the previously parameterised datasets had examined different frequency windows (from a lower bound of 0.5, 2 or 3Hz to an upper bound of 40Hz; Table 2) and we therefore parametrized the spectra across several broad and narrow frequency ranges to identify the optimal frequency windows and spatial locations for comparison across studies (see Supplementary Data). Through this process, we selected 3-40 Hz for the common pipeline, and the data was collapsed over the full scalp before being entered into the meta-analysis.

Error metrics were used to identify and exclude samples with poor model accuracy. For each data cohort, the model fit was obtained and the z-score of the model accuracy distribution was then computed to detect and exclude subjects with poor model performance. As a result, the following participants were excluded at this stage: 1 PD patient from the San Diego dataset, 2 healthy controls from the Turku dataset, 4 PD and 2 HCs from Iowa, 1 PD and 2 HCs from New Mexico, 5 PD and 14 HCs from OMEGA, and 3 PD patients from NatMEG. The number of subjects included in the final meta-analysis from each dataset is shown in Table 1. After removing outliers, the model accuracy had a mean fit of 97.04% (range = 96.54-97.8) for the PD group and 97.38% (range = 96.98-97.7) for the healthy controls.

The unadjusted and adjusted spectral plots were used to select the frequencies of interest for meta-analysis (Figure 2). We observed clear peaks within the alpha (7-12 Hz) and beta (12-30 Hz) frequency windows, but no peaks were identified within the theta or gamma bands. Thus, the peak adjusted alpha and beta power, and their respective peak frequencies, were entered into the meta-analysis, together with both aperiodic components, i.e. offset and exponent.

**Figure 2.**
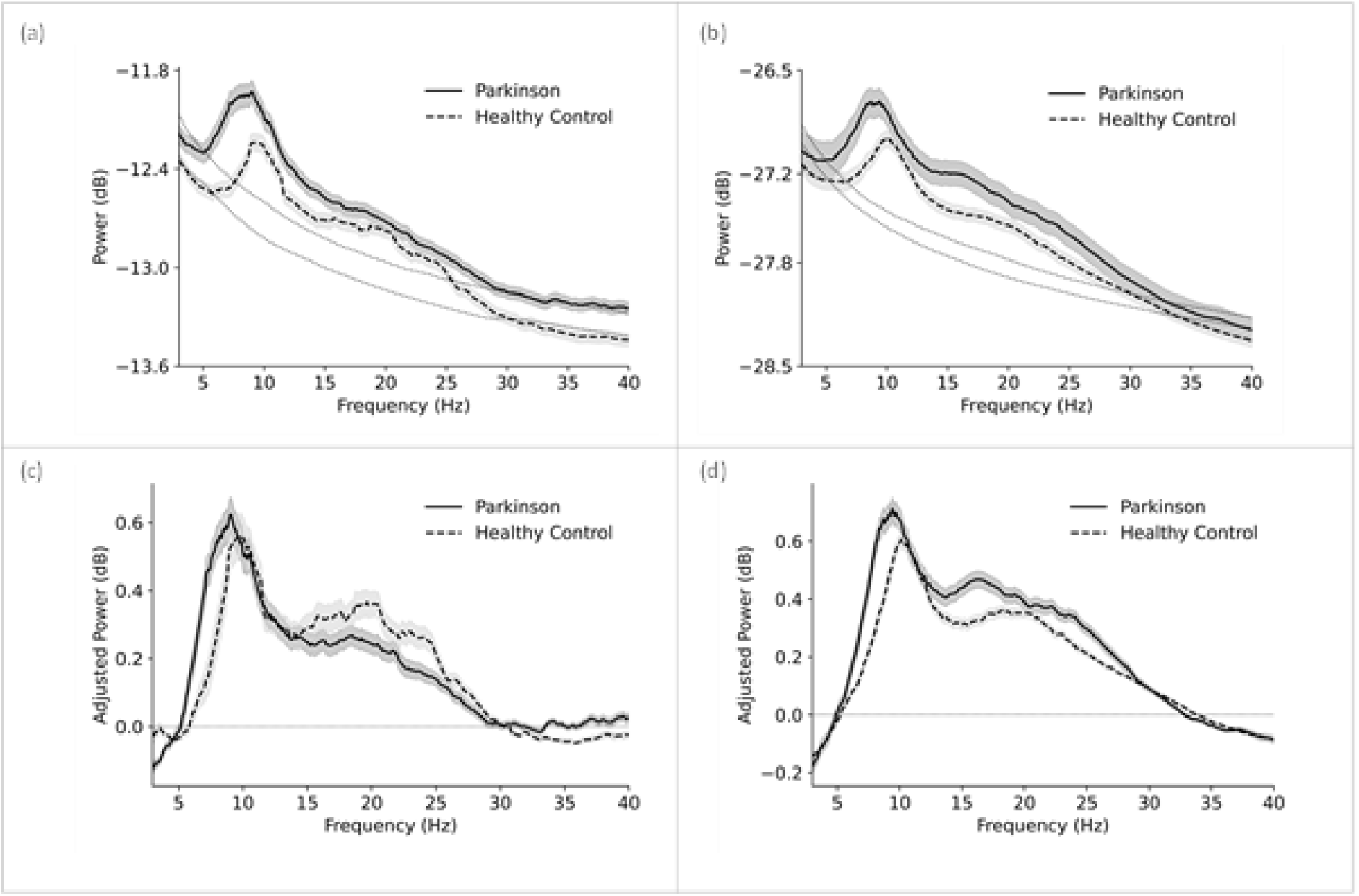
(a) Mean unadjusted power in the 2 MEG datasets. (b) Mean adjusted power (aperiodic signal subtracted) in the 2 MEG datasets (c) Mean unadjusted power in the 4EEG datasets, (d) Mean adjusted power (aperiodic signal subtracted) in the 4 EEG datasets.

### 2.4. Meta-analysis

The following features extracted from each dataset and were used in the meta-analysis: alpha power, alpha peak frequency, beta power, beta peak frequency, exponent, and offset. The *metafor* (Viechtbauer, 2007) and *robumeta* (Fisher et al., 2017) packages for R were used to perform the meta-analysis. First, the standardized mean difference and sampling variance were calculated, using the *escalc* function, between Parkinson’s patients and healthy controls, separately for each of the 6 components. Random effects models were then run for each component and mean weighted effect sizes and 95% confidence intervals were estimated using restricted maximum likelihood (REML). Heterogeneity was assessed using the Q-statistic, I^2^, R^2^ and tau^2^ indices. Forest plots were used to visualise the standardised mean difference across datasets.

## 3. Results

A total of 6 datasets involving 368 patients with Parkinson’s disease and 570 age-matched controls were included in the meta-analyses of 1) alpha peak amplitude, 2) alpha peak frequency, 3) beta peak amplitude, 4) beta peak frequency, 5) exponent, 6) offset. The mean unadjusted and aperiodic-adjusted signal is shown in Figure 2, separately for Parkinson’s patients and healthy controls, and for MEG and EEG datasets.

### 3.1. Alpha peak amplitude

A small effect of group on alpha amplitude was observed, SMD = 0.33, 95% CI = [0.15, 0.52], *p* = 0.00037, with a larger alpha amplitude for PD patients relative to controls (Figure 3A). There was limited evidence of heterogeneity across the 6 datasets: Q (5) = 6.05, *p* = 0.30, tau^2^ = 0.01, I^2^ = 22.50%.

**Figure 3.**
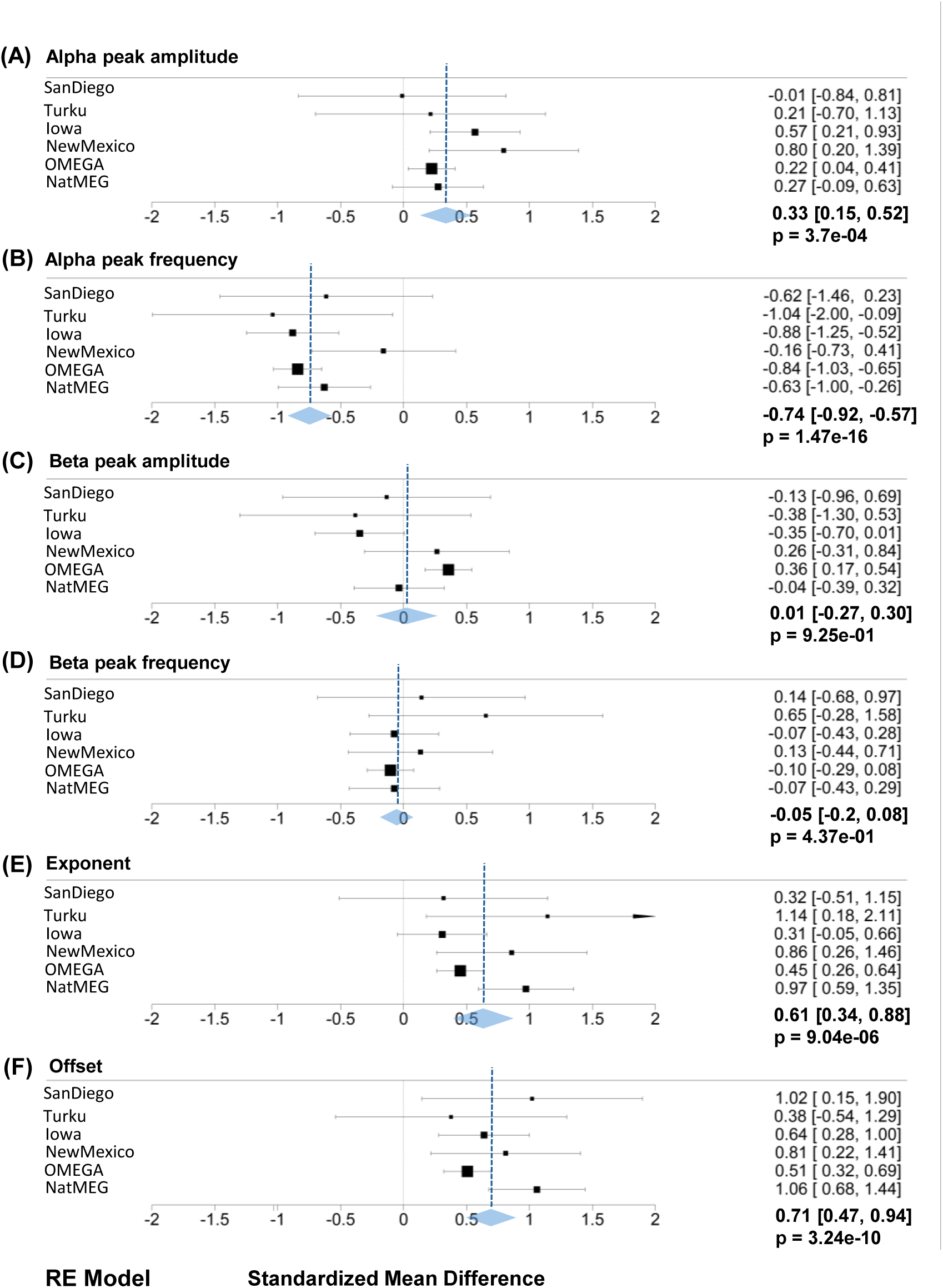
Standardized mean differences between Parkinson’s patients and healthy controls for A) Alpha peak amplitude, B) Alpha peak frequency, C) Beta peak amplitude, D) Beta peak frequency, E) Exponent and F) Offset. A positive number indicates values are higher in patients relative to controls, and a negative number indicates higher in controls relative to patients.

### 3.2. Alpha peak frequency

There was a large effect of group on alpha central frequency, SMD =-0.74, 95% CI = [-0.92,-0.57], p <0.00001, with slower alpha central frequency for PD patients relative to controls (Figure 3B). There was low evidence of heterogeneity across datasets: Q(5) = 6.24, *p* = 0.28, tau^2^ = 0.008, I^2^ = 15.76%.

### 3.3. Beta peak amplitude

There was no between-group difference in beta amplitude, SMD = 0.01, 95% CI = [-0.27, 0.30], *p* = 0.925 (Figure 3C). There was evidence of heterogeneity across datasets: Q(5) = 15.18, p = 0.009, tau^2^ = 0.06, I^2^ = 61.98% and visual inspection of the forest plot identified the OMEGA dataset as an outlier relative to the other 5 datasets. Since no effect of group was found when collapsing the data across all scalp electrodes, we performed separate random effects analyses for the frontal, central and posterior scalp regions. None of these models identified differences in beta amplitude between PD patients and controls (Frontal: SMD = 0.06, 95% CI = [-0.26, 0.39], *p* = 0.69; Central: SMD = 0.09, 95% CI = [-0.19, 0.37], *p* = 0.52; Posterior: SMD =-0.66, 95% CI = [-1.78, 0.45], *p* = 0.24).

### 3.4. Beta peak frequency

We identified no difference between PD patients and controls for beta central frequency, SMD =-0.05, 95% CI [-0.20, 0.08], p = 4.37e-01 (Figure 3D). There was no evidence of heterogeneity: Q(5) = 3.15, *p* = 0.63, tau^2^ = 0.0, I^2^ = 0.0%.

### 3.5. Exponent

PD patients had a markedly larger exponent compared to healthy controls: SMD = 0.61, 95% CI = [0.34, 0.88], *p* < 0.00001 (Figure 3E), although there was evidence of heterogeneity across datasets: Q(5) = 10.40, *p* = 0.06, tau^2^ = 0.05, I^2^ = 55.47%.

### 3.6. Offset

PD patients had a larger offset compared to controls: SMD = 0.71, 95% CI = [0.47, 0.94], *p* < 0.00001, (Figure 3F) and there was some heterogeneity across datasets: Q(5) = 7.91, *p* = 0.16, tau^2^ = 0.03, I^2^ = 41.87%.

## 4. Discussion

We conducted an advanced re-analysis and meta-analysis of resting-state M/EEG datasets to compare periodic and aperiodic components in patients with Parkinson’s disease and age-matched healthy controls. Using a common analytical pipeline, aggregating multiple, raw, open-access M/EEG datasets, we compared the power and peak frequency of the aperiodic-adjusted alpha and beta oscillations, and the aperiodic components, i.e. exponent and offset across the two groups. A total of 6 datasets were included (4 recorded using EEG and 2 using MEG), involving 938 participants (368 with Parkinson’s disease and 570 healthy controls). In summary, we identified higher alpha power and slower peak alpha frequency, in addition to higher aperiodic offset and exponent, in Parkinson’s patients relative to controls, but there were no between-group differences in beta power or beta peak frequency in this resting state data.

### 4.1. Higher peak alpha power and slower peak alpha frequency in Parkinson’s disease

The meta-analyses of alpha band measures were broadly consistent across the six included datasets, with statistical indices suggesting minimal between-study heterogeneity for both higher peak alpha power and slower peak alpha frequency in PD patients relative to controls (Sinanovic et al., 2005; Soikkeli et al., 1991; Stoffers et al., 2007; Ye et al., 2022). However, the relevance of this change in the clinical population is not immediately clear. In healthy adults, alpha rhythms are proposed to reflect decreased excitability of the underlying cortex and are modulated by both tonic and selective attention (Jensen, 2024; Thut et al. 2006). For example, increases in alpha power are observed during fatigued states (Hanzal et al. 2024; Tran et al. 2020), and linear increases in alpha activity are commonly recorded during the course of experimental sessions, as participants experience fatigue as a result of extended engagement with perceptual tasks (the “time-on-task” effect) (Benwell et al. 2019; Kasten et al. 2016).

Similarly, Parkinson’s disease presents not only with motor symptoms but also with a range of non-motor symptoms, such as fatigue and apathy (Dujardin et al. 2007; Marin 1991; Pavese et al. 2010; Sáez-Francàs et al. 2013a; Siciliano et al. 2018a; Zuo et al. 2016). Around half of PD patients experience fatigue, both in terms of reduced physical and mental energy and increased effort in cognition and perception, at higher levels than healthy individuals (Siciliano et al. 2018b). Apathy is characterised by diminished motivation, goal-directed behaviour, and emotional engagement, manifesting as indifference, lack of initiative, and impaired decision-making (Ineichen and Baumann-Vogel 2021; Sáez-Francàs et al. 2013b). Both apathy and fatigue can appear early in PD, often preceding motor symptoms, and their prevalence often increases as the disease progresses (Ineichen and Baumann-Vogel, 2021; Siciliano et al. 2020; Siciliano et al. 2018b).

Although we identified a consistent increase in alpha power in patients relative to controls, which may be related to transient or chronic episodes of fatigue and apathy, none of the datasets included in the meta-analyses included clinical fatigue measurements to test this hypothesis directly. The OMEGA dataset did include scores from the Unified Parkinson’s Disease Rating Scale Part I (UPDRS-I), which contains sub-item UPDRS-I.13, assessing fatigue over the past few weeks. However, these scores were unavailable in other datasets, precluding a formal meta-regression. Furthermore, although sub-item UPDRS-I.13 assesses fatigue, it may not serve as a reliable indicator of the fatigue state experienced at the time of M/EEG recording. As such, we cannot determine whether the observed alterations in alpha oscillations are driven by underlying pathology, cognitive changes, elevated apathy or fatigue levels, or a combination of these factors. We recommend that future research includes both physical and cognitive fatigue measures taken at the time of recording, alongside longer-term assessments, to better understand the relationship between Parkinson’s disease and increased alpha power.

In addition to increased alpha power, we also identified a slowing of peak alpha frequency in Parkinson’s patients relative to controls. Although subtle, this effect was consistent across datasets, with a mean slowing of 0.92 Hz (SD = 0.062). Individual alpha frequency gradually changes non-linearly throughout the lifespan (Aurlien et al. 2004; Cellier et al. 2021; Chiang et al. 2011; Cragg et al. n.d.; Duffy et al. 1984; Freschl et al. 2022; Grandy et al. 2013; Klimesch 1999; Knyazeva et al. 2018; Marshall et al. 2002; Turner et al. 2023) typically peaking around 10 Hz on average in young adulthood (Turner et al. 2023) followed by a gradual slowing into older age. The functional significance of individual alpha frequency (IAF) is still under debate (see Samaha & Romei, 2024, for a review) but has been proposed to represent the “sampling rate” of the perceptual system (Samaha and Postle 2015), suggesting that slowing of IAF in Parkinson’s patients may reflect disrupted perceptual or cognitive processing (Chunharas et al., 2022). Specifically, PD patients with more pronounced alpha slowing show increased disease progression and cognitive decline in longitudinal studies (Olde Dubbelink et al., 2013). These findings indicate that pathological changes in neural competence in Parkinson’s disease may directly impact neural oscillations. However, these changes could also reflect secondary effects of non-motor symptoms commonly observed in PD.

Finally, we evaluated the consistency of our re-analysis results with those reported in the original publications. Aperiodic-adjusted alpha power had previously been reported for three of the included datasets (i.e. OMEGA, NatMEG and New Mexico). Since we re-analysed all datasets using a common processing pipeline, we compared our outputs to the results from the original articles. Our re-analysis was consistent across these datasets: all 3 papers had previously reported an increased alpha power in PD patients and we replicated this effect. For the remaining three datasets (i.e. San Diego, Turku, and Iowa), which had not previously been analysed using spectral parameterisation, we replicated the absence of group-level differences originally described in the San Diego dataset. However, the findings from our re-analysis differed from the reported results of the Turku and Iowa studies.

Of the six datasets included in our analyses, only one (NatMEG) had previously examined peak alpha frequency between groups and reported no differences. In contrast, our re-analysis of the raw data identified a significant slowing of peak alpha frequency in Parkinson’s disease relative to controls in the NatMEG dataset (and consistently across all datasets). This discrepancy is likely due to differences in the selection of electrodes analysed; the original analysis focused on a fronto-central sensorimotor region of interest, whereas our approach used full-scalp data, including the posterior electrodes. These findings underscore the impact of analytical choices, such as electrode selection and the use of aperiodic-adjusted versus unadjusted spectral analysis, on the outcomes of M/EEG data analysis. To improve transparency, reproducibility, and comparability across studies, we recommend that researchers report both adjusted and unadjusted power spectra, and, where possible, make raw data and analysis scripts publicly available or on request.

### 4.2. No group-level differences in resting-state beta activity

We found no differences in resting-state beta power or beta peak frequency between individuals with Parkinson’s disease and healthy controls. This is perhaps unsurprising, given that we analysed resting state activity, whereas beta band activity (13-30 Hz) is more strongly associated with motor preparation and execution (Alayrangues et al. 2019; Rhodes et al. 2018). Our re-analysis was consistent with the original findings from the NatMEG, New Mexico and San Diego datasets, all of which reported no between-group differences in beta power. In contrast, the original analyses of the OMEGA, Iowa, and Turku datasets reported reduced beta power in Parkinson’s patients. However, our re-analysis identified no group-level differences in the Iowa and Turku datasets (which had not been parameterised), and higher beta power in PD patients in the (parameterised) OMEGA dataset.

Our decision to analyse peak amplitudes within canonical frequency bands may have been sensitive enough to capture important pathological electrophysiological changes in Parkinson’s disease. Beta activity often occurs in transient bursts with varying duration and amplitude (He et al. 2020; Szul et al. 2023; Tinkhauser, Pogosyan, Little, et al. 2017b). Notably, beta bursts occurring close in time to motor movements are associated with slower response times (Little et al. 2019), and neurofeedback-driven suppression of these bursts leads to faster reaction times (He et al. 2020). Thus, analysing the temporal dynamics of beta bursts may offer a more sensitive and reliable method for investigating cortical motor impairments in PD.

While heterogeneity across the six datasets was low for peak beta frequency, there was high heterogeneity for beta power. The OMEGA dataset emerged as an outlier, with PD patients showing significantly larger beta amplitudes than controls. This difference may reflect greater symptom severity of the patients in this cohort, as indicated by higher UPDRS-III and Hoehn & Yahr scores, consistent with prior findings linking increased beta power to poor motor function (Wiesman, Vinding, et al. 2025). Additionally, the PD patients in the OMEGA dataset were substantially older than the controls (mean age PD = 65, SD = 9.1; mean age controls = 44.4, SD = 20.8) and elevated beta activity is commonly observed in older adults (Park et al. 2025; Xifra-Porxas et al. 2019). It is important to highlight that the version of the OMEGA dataset used in the original report (Wiesman et al. 2023a) included a smaller sample of control participants, with closer age matching between the patient and control groups. In contrast, in the current study we used a larger subset of the OMEGA data, which includes a broader age range in the control group. This difference in sample composition may contribute to the discrepancies in beta power observed between the two studies. In short, our results suggest that *resting-state* beta power and peak frequency are unlikely to serve as reliable electrophysiological biomarkers of Parkinson’s disease, and further synthesis of active-state M/EEG should be conducted.

### 4.3. Higher aperiodic exponent and offset in Parkinson’s disease

Compared to healthy controls, Parkinson’s patients had a consistently larger aperiodic exponent and offset across the six datasets. The functional relevance of the M/EEG exponent has been variously interpreted as an index of the neural excitation-inhibition ratio (Darmani et al. 2023; Helson et al. 2023; Wiest et al. 2023), oscillatory slowing (Wiesman et al. 2023b), and oscillation damping (Evertz et al. 2022). We also observed a large effect of group on the aperiodic offset, with PD patients showing larger offset values than healthy controls. Taken together, these aperiodic changes could indicate alterations in excitation-inhibition balance and increased baseline neural activity, reflecting pathological changes in network dynamics associated with Parkinson’s disease. Regardless of the physiological interpretation, our results suggest that elevated aperiodic exponent and offset are likely to be robust and reproducible characteristics across PD studies (Bush et al. 2024; Clark et al. 2023; da Silva Castanheira et al. 2024; Darmani et al. 2023; Liu et al. 2024; Martin et al. 2018; Monchy et al. 2023; Mostile et al. 2019; Rosenblum et al. 2023; Vinding et al. 2024a; Wang et al. 2022b; Wiest et al. 2023; Zhang et al. 2022b). Additionally, the aperiodic offset and exponent may also be potential indicators to differentiate early-from late-onset PD (Wu et al. 2023).

Three of the included datasets (New Mexico, NatMEG, and OMEGA) had been analysed using spectral parameterisation. These studies reported elevated aperiodic offset and exponent values in PD patients, which our meta-analysis confirms are reliable markers consistently found in Parkinson’s disease across the datasets. On a methodological note, although our analysis pipeline is broadly comparable, each study employed slightly different parameterisation approaches, with minor variations in the parameters and frequency ranges. A systematic investigation of how these methodological choices influence the spectral parameterisation outputs, and the establishment of best-practice guidelines would improve consistency across research groups. Reliable measures are critical for meaningful comparisons across individuals and groups over time, indicating that observed changes reflect biological factors rather than measurement errors (Ding et al. 2022; Ip et al. 2018). Both aperiodic offset and exponent demonstrated good test-retest reliability (McKeown, Finley, et al. 2024), suggesting these measures are stable over time and capable of capturing individual differences in neural activity. This reliability positions them as promising markers for investigating neural mechanisms and clinical conditions.

Whilst our results, and previous studies (McKeown, Jones, et al. 2024b; Vinding et al. 2024a) provide evidence that the aperiodic offset can distinguish PD patients from healthy controls, its functional relevance remains unclear. Although the aperiodic offset and exponent have been associated with motor symptom severity in PD (Clark et al. 2023), conflicting evidence has also been reported (Bush et al. 2024). Similarly, cognitive decline in PD also has been associated with aperiodic activity in some studies (Rosenblum et al. 2023; Wiesman et al. 2023a), but this association could not be replicated in others (Bernasconi et al. 2023a). Overall, it remains uncertain whether the aperiodic offset and exponent are directly linked to cognitive and motor symptoms in PD. Given the contradictory evidence, further research is needed to clarify these relationships.

### 4.4. Limitations and future directions

In our re-analysis of existing raw M/EEG datasets, we developed a common processing pipeline that was applied to all 6 datasets, to ensure that the aperiodic and periodic components were extracted from consistently cleaned and processed signals. As shown in the Supplementary data, varying the frequency ranges used for spectral parameterisation substantially affects the outputs, particularly the estimate of the aperiodic exponent. For example, parameterising the spectrum from 1-20 Hz results in a smaller exponent compared to using 3-40 Hz. In addition, identifying peaks within the canonical frequency bands involves researcher-led definitions of the upper and lower limits of those bands of interest (e.g., alpha may be variously defined as 8-12Hz or 8-13Hz, or further subdivided into low (8-10Hz) and high (10-12Hz) alpha). Although both the clinical and control groups in this study were processed using the same pipeline, allowing for a comparison of standardized mean differences between the two groups, the impact of these experimenter-defined choices on the outputs of spectral parameterisation is under-explored.

Second, to facilitate data synthesis, we chose to analyse only scalp-wide averages of the data across all available electrodes, rather than conducting region of interest analysis, or an analysis of individual electrodes. While this approach ensured comparability across datasets, it likely obscured any subtle, region-specific changes in periodic and aperiodic activity (Donoghue 2024). Given that the six datasets differed in the number and placement of available electrodes, it was deemed not possible to perform a direct comparison to such a degree of sensitivity. However, we conducted a preliminary pilot analysis comparing aperiodic exponents across frontal, central and posterior regions found no substantial regional differences between PD patients and healthy controls (see Supplementary data). This absence of regional variation may reflect the task-free nature of the resting-state recordings used in this study and it is possible that task engagement modulates aperiodic activity differently across brain regions (Gyurkovics et al., 2022).

Our decision to focus the analysis on alpha and beta frequency windows was data-driven, based on the presence of observed peaks in the grand-average spectral plots (Figure 2). Consequently, we did not perform a synthesis of other frequency bands, such as the theta band, which has been reported to change in Parkinson’s disease and may be a biomarker of PD-related cognitive decline (Zawiślak-Fornagiel et al., 2023; Ye et al., 2022; Iyer et al., 2020). We also chose to only include PD patients who were tested ON-medication, due to the low availability of both ON- and OFF-medication datasets. It remains possible that different effects, particularly in beta power and aperiodic components, would emerge when comparing patients tested OFF medication states to controls. Further research examining ON versus OFF medication states, as well as recording during active movement, is needed to evaluate the reliability of spectral parameterisation as a biomarker for PD. Additionally, it remains unclear whether the periodic and aperiodic changes observed in Parkinson’s disease represent specific disease biomarkers or simply correlate with disease presence.

## 5. Conclusion

Parkinson’s patients exhibit increased alpha power and slowed peak alpha frequency, alongside elevated aperiodic offset and exponent values during resting states. This meta-analysis highlights the potential of M/EEG as a tool for identifying consistent cortical electrophysiological changes in Parkinson’s disease. Further research is needed to refine these electrophysiological measures (particularly with best-practice guidelines for parameter selection during the processing stages), and to explore their potential clinical applications in the diagnosis and treatment of neurodegenerative conditions.

## Acknowledgements

HN is supported by a University of Stirling Institute for Advanced Studies PhD scholarship. The funder had no role in the design of the study, data analysis, or preparation of the manuscript. We would like to express our sincere gratitude to all individuals and institutions who contributed to this research. In particular, we thank the Quebec Parkinson’s Network at McGill University, the Swedish National Facility for Magnetoencephalography Parkinson’s Disease Dataset at the Department of Clinical Neuroscience, Karolinska Institute, Professor James Cavanagh (University of New Mexico), Professor Nandakumar Narayanan (University of Iowa), Dr. Henry Railo (University of Turku), and Dr. Alexander Rockhill (University of Oregon) for generously providing access to their data.

## Notes

### Competing Interest Statement

The authors have declared no competing interest.

