## Supplementary Data for "A meta-analysis of periodic and aperiodic M/EEG components in Parkinson’s disease"

### 1. Rationale for selecting the 3-40 Hz range for spectral parameterisation

A recent study (Donoghue, 2024) reported that approximately 50% of previous research decomposing periodic and aperiodic components and estimated exponent values used a frequency range below 43 Hz. Furthermore, 85% of studies employing frequency ranges below 40 Hz selected ranges with a minimum frequency of 3 Hz. While some research recommends conducting spectral parameterisation within a narrow frequency window, others argue that broader frequency windows improve model performance and affect key parameters such as the exponent and offset (Ameen et al. 2024). To systematically assess these effects, we tested multiple window sizes during the development of our spectral parameterisation pipeline, specifically: 1–20 Hz, 1–30 Hz, 1–40 Hz, 1–50 Hz, 1–60 Hz, 1–70 Hz, 1–80 Hz, 1–90 Hz, 3–20 Hz, 3–30 Hz, 3–40 Hz, 3–50 Hz, 3–60 Hz, 3–70 Hz, 3–80 Hz, 3–90 Hz). The results are shown in Figure S.1 below.

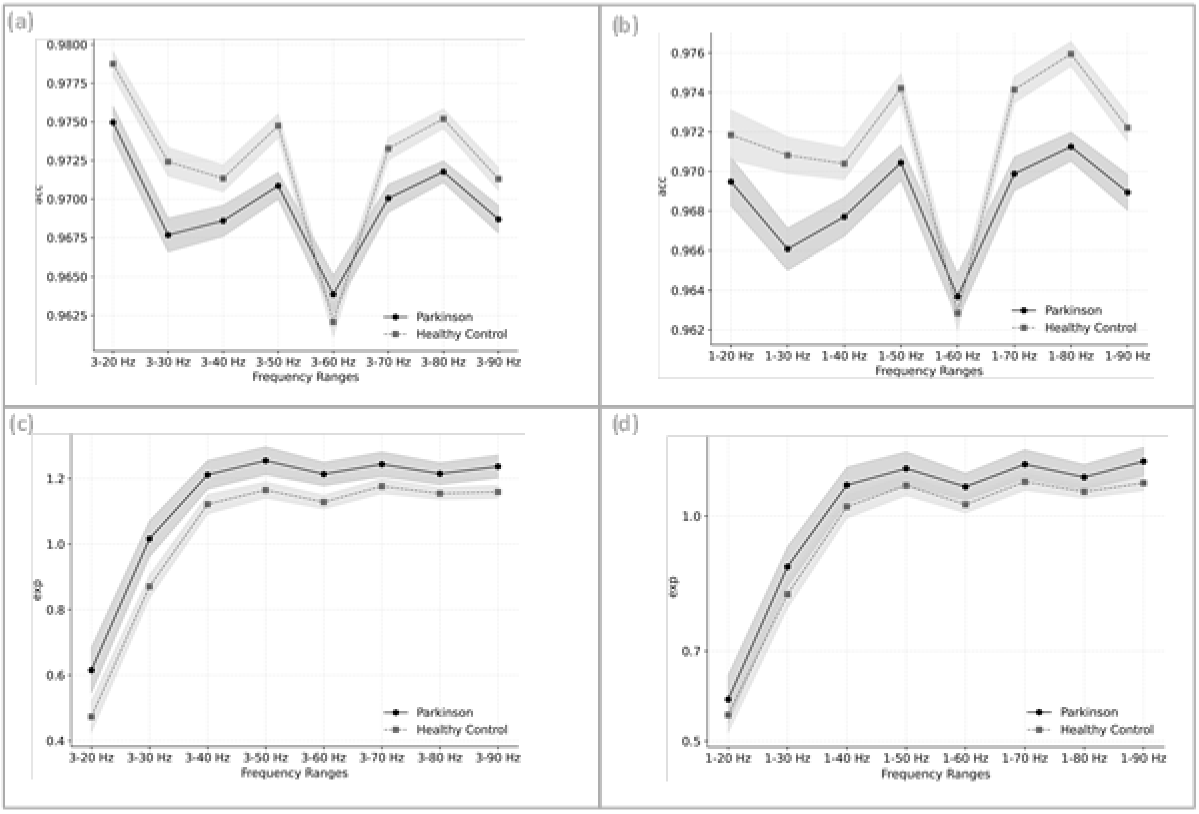


**Figure S.1.** Effects of different frequency ranges on *specparam* model variables across all 6 datasets (A) Impact of [3- F] frequency window on *specparam* model accuracy. (B) Impact of [1-F] frequency window on *specparam* model accuracy. (C) Effect of [3- F] frequency range on exponent variation. (D) Effect of [1- F] frequency range on exponent variation. Note: Each point represents mean and 95% confidence intervals of the variable.

Figures S.1A and S.1B illustrate the parametrization of spectral accuracy across different frequency ranges. windows had a high accuracy of > .96. A slight decrease in model fit was observed in the ranges of 1-60 Hz and 3-60 Hz due to power line noise. We also found that frequency ranges 1–20 Hz, 3–20 Hz, 1–30 Hz, and 3–30 Hz did not provide stable exponent values, with smaller windows generating shallower slopes, although exponent values plateaued in broader windows over 40 Hz (Figures S.1C and S.1D). Based on these observations, and to maintain consistency with prior research, we selected the 3–40 Hz range for our analysis.

### 2. Rationale for analysing the full scalp rather than regions of interest

It has been suggested that parameterization should be performed within specific scalp regions, as averaging power spectra across the entire scalp may obscure region-specific electrophysiological changes (Donoghue 2024). To investigate this, we applied our analysis pipeline to M/EEG signals from electrodes grouped into Frontal, Central, and Occipital regions, as well as an averaged power spectrum signal across all electrodes, referred to as the AllScalp region.

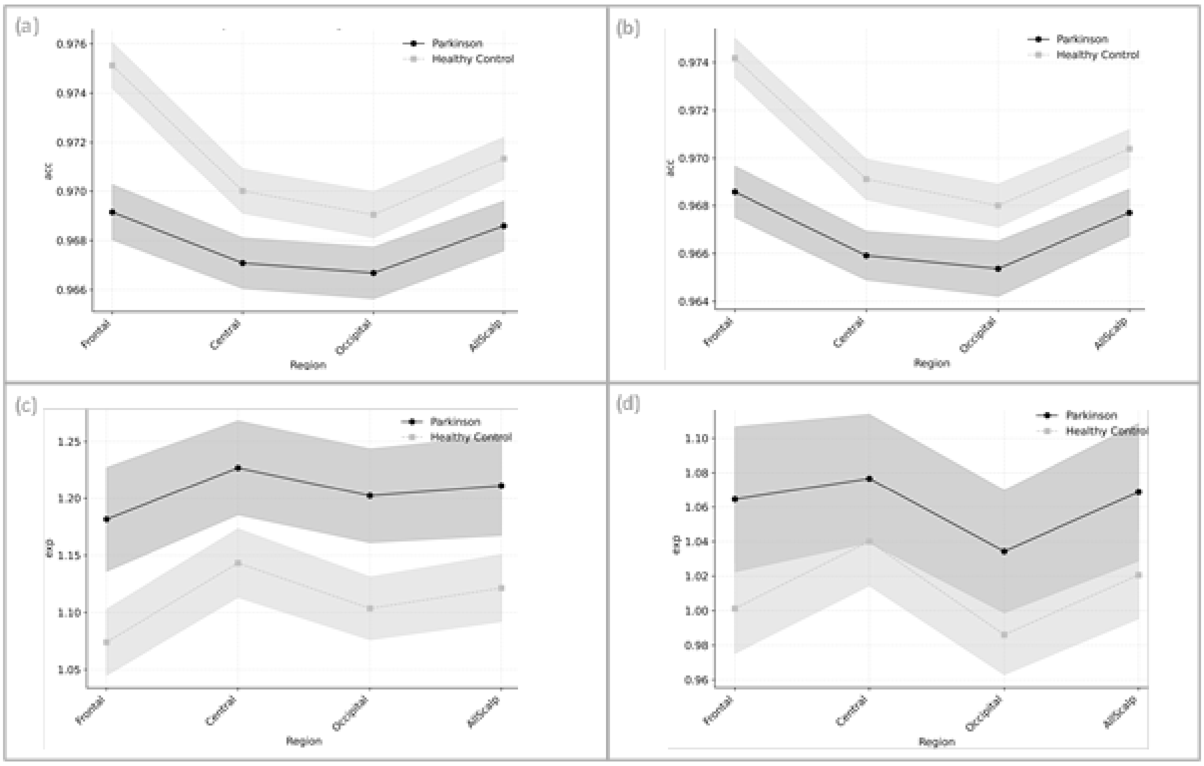


**Figure S.2** Effect of different regions on (FOOOF) model variables across all 6 datasets (A) Impact of regions on (FOOOF) model accuracy in [3 - 40] Hz frequency window. (B) Impact of regions on (FOOOF) model accuracy in [1 - 40] Hz frequency window (C) Effect of regions on exponent variation in [3 – 40] Hz frequency range. (D) Effect of regions on exponent variation in [1 - 40] Hz frequency range. Note: Each point represents mean and 95% confidence intervals of the variable.

Figures S.2A and S.2B illustrate accuracy across different regions in ranges 3-40 Hz and 1-40 Hz respectively. All regions of interest showed high model accuracy of >.96. the difference in model fit between the two groups is consistent. Figure S.2C and S.2D shows that the between-group difference in exponent is relatively consistent across the regions of interest (shown for analysis in the range of [3-40] Hz and [1-40] Hz respectively).

Additionally, due to variations in recording modalities across the recruited datasets (i.e., EEG and MEG), the absence of individual MRI data or head models for precise electrode localization, differences in the number of channels, and the use of different EEG electrode placement systems, we used the averaged power spectrum signal across all electrodes to minimize the risk of incorrect electrode selection for each region. Overall, based on our observations regarding the minimal impact of regional differences on exponent estimation and model accuracy (figure S.2C and S.2D), as well as the limitations in electrode position mapping, utilizing the averaged power spectrum signal for decomposing aperiodic and periodic components was deemed an appropriate approach.

*Table S.1 Mean and standard deviation of targeted features for PD patients.*

| DATASET | PD exponent | PD offset | PD alpha-amplitude | PD alpha-Peak Frequency | PD beta-amplitude | PD beta-Peak Frequency |
|---|---|---|---|---|---|---|
| SanDiego (13 PD$^{U}$) | 1.24 (0.14) | -11.573 (0.241) | 0.605 (0.341) | 9.11 (1.125) | 0.398 (0.122) | 19.804 (3.539) |
| Turku (6 PD$^{U}$) | 1.236 (0.16) | -11.34 (0.222) | 0.522 (0.218) | 8.567 (1.547) | 0.261 (0.3) | 19.45 (3.797) |
| Iowa (115 PD$^{U}$) | 1.036 (0.341) | -11.59 (0.585) | 0.754 (0.294) | 8.652 (1.506) | 0.3 (0.184) | 17.403 (3.556) |
| NewMexico (24 PD$^{U}$) | 1.138 (0.236) | -11.56 (0.464) | 0.746 (0.221) | 9.367 (1.536) | 0.435 (0.166) | 18.05 (3.017) |
| OMEGA (154 PD$^{U}$) | 1.373 (0.518) | -25.81 (0.998) | 0.701 (0.265) | 9.13 (1.369) | 0.475 (0.193) | 18.503 (3.7) |
| NatMEG (56 PD$^{U}$) | 1.148 (0.254) | -27.14 (0.341) | 0.735 (0.253) | 9.036 (1.175) | 0.453 (0.147) | 16.542 (3.784) |
| **Mean (SD)** | **1.195 (0.274)** | **EEG: -11.516 (0.378), MEG: -26.475 (0.67)** | **0.677 (0.265)** | **8.977 (1.376)** | **0.387 (0.185)** | **18.291 (3.565)** |

*Note:U Number of PD patients in each dataset. EEG datasets: SanDiego, Turku, Iowa, NewMexico. MEG datasets: OMEGA, NatMEG.*

*Table S.2 Mean and standard deviation of targeted features for HC individuals.*

| DATASET | HC exponent | HC offset | HC alpha-amplitude | HC alpha-Peak Frequency | HC beta-amplitude | HC beta-Peak Frequency |
|---|---|---|---|---|---|---|
| SanDiego (10 HC[a]) | 1.197 (0.117) | -11.862 (0.31) | 0.609 (0.364) | 9.845 (1.186) | 0.419 (0.182) | 19.275 (3.669) |
| Turku (42 HC[a]) | 1.02 (0.187) | -11.663 (0.918) | 0.454 (0.328) | 10.006 (1.279) | 0.35 (0.2) | 18.247 (0.471) |
| Iowa (42 HC[a]) | 0.937 (0.258) | -11.948 (0.465) | 0.575 (0.364) | 9.94 (1.298) | 0.366 (0.201) | 17.657 (3.041) |
| NewMexico (23 HC[a]) | 0.943 (0.209) | -11.907 (0.37) | 0.542 (0.281) | 9.639 (1.802) | 0.393 (0.145) | 17.634 (3.103) |
| OMEGA (411 HC[a]) | 1.183 (0.379) | -26.249 (0.804) | 0.639 (0.283) | 10.206 (1.24) | 0.409 (0.182) | 18.909 (3.957) |
| NatMEG (64 HC[a]) | 0.932 (0.187) | -27.48 (0.298) | 0.66 (0.288) | 9.75 (1.08) | 0.459 (0.18) | 16.79 (3.025) |
| **Mean (SD)** | **1.035 (0.223)** | **EEG: -11.845 (0.516), MEG = -26.865 (0.551)** | **0.58 (0.318)** | **9.898 (1.314)** | **0.399 (0.182)** | **18.085 (2.878)** |

*Note: ª Number of HC participants in each dataset. EEG datasets: SanDiego, Turku, Iowa, NewMexico. MEG datasets: OMEGA, NatMEG.*

*Table S3. Mean and standard deviation of frontal beta amplitude for PD and HC groups.*

| DATASET | PD beta amplitude | HC beta amplitude |
|---|---|---|
| San Diego | 0.362 (0.109) | 0.412 (0.195) |
| Turku | 0.268 (0.26) | 0.339 (0.168) |
| Iowa | 0.325 (0.181) | 0.368 (0.191) |
| New Mexico | 0.456 (0.178) | 0.432 (0.158) |
| OMEGA | 0.405 (0.199) | 0.302 (0.165) |
| NatMEG | 0.49 (0.151) | 0.481 (0.191) |
| **Mean (SD)** | **0.384 (0.18)** | **0.389 (0.178)** |

*Table S4. Mean and standard deviation of central beta amplitude for PD and HC groups.*

| DATASET | PD beta amplitude | HC beta amplitude |
|---|---|---|
| San Diego | 0.446 (0.155) | 0.424 (0.191) |
| Turku | 0.266 (0.3) | 0.326 (0.213) |
| Iowa | 0.385 (0.402) | 0.447 (0.042) |
| New Mexico | 0.418 (0.154) | 0.428 (0.148) |
| OMEGA | 0.523 (0.195) | 0.426 (0.197) |
| NatMEG | 0.477 (0.143) | 0.469 (0.178) |
| **Mean (SD)** | **0.419 (0.225)** | **0.42 (0.162)** |

*Table S5. Mean and standard deviation of occipital beta amplitude for PD and HC groups.*

| DATASET | PD beta amplitude | HC beta amplitude |
|---|---|---|
| San Diego | 0.326 (0.142) | 0.393 (0.168) |
| Turku | 0.264 (0.297) | 0.371 (0.211) |
| Iowa | 0.249 (0.0358) | 0.379 (0.0439) |
| New Mexico | 0.45 (0.175) | 0.43 (0.168) |
| OMEGA | 0.506 (0.18) | 0.426 (0.189) |
| NatMEG | 0.443 (0.187) | 0.499 (0.201) |
| **Mean (SD)** | **0.373 (0.169)** | **0.416 (0.164)** |
